# Loss of Nuclear Deformability of Breast Cancer Cells by the Disruption of Actin Filaments

**DOI:** 10.1101/854562

**Authors:** Ezgi Antmen, Utkan Demirci, Vasif Hasirci

**Affiliations:** BIOMATEN, Middle East Technical University (METU) Center of Excellence in Biomaterials and Tissue Engineering, Ankara, Turkey; METU, Departments of Biotechnology, Ankara, Turkey; Biological Sciences, Ankara, Turkey; Acibadem Mehmet Ali Aydinlar University, Department of Medical Engineering, Atasehir, Istanbul, Turkey; Department of Radiology, School of Medicine, Stanford University, Palo Alto, CA, USA

**Keywords:** actin filaments, nuclear deformation, breast cancer, micropattern, mechanotransduction

## Abstract

It is well known that chemical and biomechanical interactions between the nucleus and cytoskeleton are involved in and critical for movement, migration and nuclear positioning of cancer cells. Through nucleo-cytoskeletal coupling, proteins of the LINC complex and the nuclear envelope are capable of transducing cytoplasmic mechanical input across the nuclear membrane; however, their functional importance in the behavior of cancer cells and their nuclei has never been directly tested. In this study, our assumption was that the differences in the malignancies of breast cancer cells are the result of the differences in their nuclear deformation and its expression can be amplified on micropatterned surfaces. Based on this, our hypothesis was that the level of completeness of polymerization of actin filaments can affect nuclear deformability, and as a result, the metastatic capability of the cancer cells. In order to prove this we disrupted the polymerization of the actin filaments by using two drugs, Cytochalasin D and Jasplakinolide, which caused impaired propagation of intracellular forces, prevented nuclear deformation and increased in the expression levels of Lamin A/C and Nesprin-2 in malignant breast cancer cells. Our findings suggest that activity of these two proteins is critical for nucleo-cytoskeletal force transmission. More importantly, actin filament disruption can prevent the distortions in nuclear morphology and as a result avoid the development of cancer metastasis.

## 1. Introduction

All cells are continuously exposed to mechanical forces created by the extracellular environment and they can sense and respond to these forces through mechanotransduction. Mechanotransduction is the process by which cells convert mechanical stimuli into biochemical signals.^1^ Mechanotransduction has a role in the regulation of blood pressure, remodeling of bone, maintenance of muscle, perception of touch and sound, cell growth, migration, gene expression, morphological changes, proliferation and differentiation as well as tissue morphogenesis and the pathogenesis of various diseases, such as cardiovascular diseases, osteoporosis and cancer.^2^ During force transmission through a cell, constant reorganization of the cell interior is needed, and this reorganization is realized by cytoskeletal remodeling and directed by mechanosensitive signaling pathways. The structural link between the cytoskeleton and nucleus play an important role in this force transmission and it is referred to as the nucleo-cytoskeleton coupling. Through this coupling, cell surface forces and substrate rigidity can change nuclear morphology and external forces can be transmitted to the nucleus and change chromosome movement and nuclear positioning. Actin filament networks of the cytoskeleton, as one of the most important components of nucleo-cytoskeleton coupling, play a critical role in maintenance of the form and orientation of nuclei, nuclear morphology and cellular stiffness through their polymerization and depolymerization.^3,4^ In cancer studies, understanding of force transmission has been gaining importance because increased actomyosin contractility can induce cell proliferation, ECM remodeling, ECM stiffening, invasion and metastasis of tumor cells.^5^ Tumor cells are known to alter their own mechanical properties and change their responses to external physical cues. Thus, the development and metastasis of cancer are closely regulated by mechanical stresses applied on the cytoskeleton and nucleus and their responses to these forces. On the other hand, nucleus is the main component of a cell which regulates the gene expression and protein synthesis.^6^ It is also known that the nucleus is the major obstacle of a cell in migration through confined openings since it is the main contributor to the total cell stiffness, it is also the largest organelle of a cell and almost an order of magnitude stiffer than the cytoplasm.^7–9^ Thus, nucleus of a metastatic or migrating cell has to be softer and must be able to deform more than the nuclei of the other cells. Supporting this, it was reported that knockdown of the expression of nuclear envelop protein, lamin A, causes a decrease in nuclear stiffness and increases cell motility and ability to migrate.^10,11^

Since force transmission through a cell is important in the migration and metastasis of a cancer cell, modifications in surface features of the substrate has been widely used in in vitro cell biology studies in order to manipulate cell responses to the external forces or to give surfaces cell discrimination capabilities. Creation of substrate surfaces with topographic features is one of the most commonly used surface modification approach to study the cell-substrate interactions. The topographies either mimic sizes and shapes of features found in the natural environment of cells or they have artificial geometries designed for specific reasons. Since the size of cells, adhesion proteins and their ligands are in the nanometer to micrometer scale, modifications at this level are required on the substrates to create the architecture that could mimic the extracellular matrix (ECM) of a cell. Common shapes used in the fabrication of surface topographies are pores, gratings, wells, pits, cones, posts, pillars, grooves or mesh-like structures that can be organized in a regular pattern.^12–16^ Several studies have shown that many cell types react strongly to micro and nano features on substrate surfaces.^13–19^ On such surfaces, changes have been reported in the cell behavior such as adhesion,^20,21^ alignment,^21–23^ morphology (cytoskeletal organization),^17,24^ proliferation,^25^ viability,^26^ gene expression and differentiation.^14,27,28^

In this study, main hypothesis was that the difference in the stiffness of the nuclei of malignant and benign breast cancer cells can be used as a discriminating factor in cancer detection because of the substantial deformation of the nuclei of the malignant cells that could be quantified through the extent of deformation of the cell nuclei. Although it has been reported that biomechanical properties of the cell and nucleus give information about the state of cancer progression,^29,30^ the interaction between the components of mechanotransduction complex (LINC; linkers of nucleoskeleton and cytoskeleton proteins, nuclear envelope proteins, actin filaments) and deformation of a nucleus has never been directly measured. The aim of this study was to explore the relation between the proteins of mechanical force transmission, nuclear deformability and level of malignancy of a tumor cell by inhibiting the polymerization and depolymerization of actin cytoskeleton. Three breast cancer cell lines with different malignancy levels (benign MCF10A, malignant but noninvasive MCF7, malignant and highly invasive MDAMB231) were used to compare nuclear deformability and behavior (spreading, attachment) of these cells and expression of their nuclear membrane proteins on micropatterned surfaces to support our hypothesis. Cells were seeded on polymethyl methacrylate (PMMA) surfaces decorated with square prism pillars (4×4 μm^2^ surface area, 4 μm gap size, and 8 μm height). Two different drugs, selective for actin and inhibit its polymerization, were used: Cytochalasin D (inhibits actin polymerization by capping filaments that bind to monomers) and Jasplakinolide (prevents actin depolymerization, forms actin aggregates and blocks focal adhesion kinase (FAK) signaling). Also, the relation between the deformability of nucleus, inhibition of actin polymerization and the expression of Lamin A/C and Nesprin-2 were investigated.

## 2. Experimental Section

### 2.1. Preparation of Micropatterned PMMA Films

Micropillar arrays were designed at METU BIOMATEN and outsourced to be fabricated using photolithography.^13,15,16^ The micropillars were in square prism shape, 8 μm tall with 4×4 μm^2^ area and 4 μm interpillar gap. Using wafers as a template, polydimethylsiloxane (PDMS) molds were prepared with Sylgard 184 silicone prepolymer and Sylgard 184 curing agent (Dow Corning Company, UK) (10:1 w/w) and cured for 4 h at 70 °C. PMMA films (Mw~996 kDa) (Sigma, Germany) were made by solvent removal from a solution (10% w/v in chloroform) on the PDMS mold which was air dried for 12 h at room temperature. As a control, smooth PDMS molds were used to obtain smooth PMMA films.

### 2.2. Characterization of PMMA Films

After coating with Au/Pd under vacuum, surface patterns of the PMMA films were examined with SEM (400F Field Emission SEM, USA).

### 2.3. *In Vitro* Studies

#### 2.3.1. Culture of Breast Cancer Cell Lines

MCF-10A cells were cultured in DMEM/F12 medium (Sigma, USA) supplemented with 5% fetal bovine serum (FBS) (Lonza, USA), EGF 20 ng/mL (Sigma), insulin 10 μg/mL (Sigma), hydrocortisone 0.5 mg/mL (Sigma), cholera toxin 100 ng/mL (Sigma) and 100 units/mL penicillin (Sigma, USA). MCF-7 cells were cultured in DMEM low glucose medium (Lonza, USA) supplemented with 10% fetal bovine serum (FBS) (Lonza, USA), 100 U/mL penicillin (Sigma, USA). MDA-MB-231 cells were cultured in DMEM high glucose medium (Lonza, USA) supplemented with 10% fetal bovine serum (FBS) (Lonza, USA), 100 U/mL penicillin (Sigma, USA).

PMMA films were sterilized by UV for 15 min. Cells were seeded at a density of 5×10^4^ cells/film in 100 μL of their specific growth medium, left for 6 h for adhesion at 37 °C, 5% CO_2_. Tissue culture plates (TCPS) and smooth (S) PMMA films were used as controls.

#### 2.3.2. Testing of Actin Inhibitor Drugs

##### 2.3.2.1. Dose Optimization with Alamar Blue Assay

Two different drugs (Cytochalasin D and Jasplakinolide) were used and solutions of both drugs in dimethyl sulfoxide (DMSO) were prepared. Final concentrations of both drugs in 1 mL of culture medium were in the range 0.01-10 μM for Cytochalasin D and 0.01-5 μM for Jasplakinolide. Control samples without drugs were prepared with the same highest amount of DMSO for both drugs. Cell were cultured for 24 h (day 1) in tissue culture plate (TCPS) and on micropatterned films (P), and then drugs were used on the cells for another 24 h (day 2). The half maximal inhibitory concentration (IC50) of Cytochalasin D and Jasplakinolide on the cell viability before and after drug treatment was determined by Alamar Blue assay. Effect of these chemicals on prevention of nuclear deformation of three cell lines was observed and quantified under fluorescence microscopy.

### 2.4. Microscopy Studies

#### 2.4.1. Fluorescence Microscopy

Cell seeded samples were fixed in 4% paraformaldehyde, permeabilized with 1% Triton-X 100 solution (Applichem, Germany) and incubated in BSA blocking solution (1%, w/v, in PBS) at 37 °C for 30 min. Then, films were incubated in Alexa Fluor 488® labeled Phalloidin solution (1:50, in 0.1% BSA in PBS) for 1 h, 37 °C, and then DAPI (1:1000, in 0.1% BSA in PBS) (Invitrogen, USA) was used for 5 min at room temperature to stain the nuclei. Fluorescence micrographs were obtained with an upright fluorescence microscope (Zeiss Axio Imager M2, Germany). The micrographs were obtained by using black and white filter of the microscope and pseudocolored with green (actin) and red (nucleus) by the software system of the microscope.

#### 2.4.2. Confocal Microscopy for Immunocytochemistry (ICC)

Confocal micrographs of the cells were obtained using an upright Confocal Laser Scanning Microscope (CLSM) using 488 nm, 532 nm, and 630 nm lasers (Leica SPE, Germany). Samples were fixed, permeabilized and blocked in the same way in the previous section. Then, they were incubated for 1 h at 37 °C with Alexa Fluor 532 Phalloidin (Invitrogen, USA) to stain the actin cytoskeleton. For Lamin A/C and Nesprin-2, the antibodies specific to these proteins (Anti-Nesprin-2 ab57397 and anti-Lamin-A ab8980, Abcam, UK) were used according to manufacturer’s directions. TOPRO-3 (1:1000, in PBS) (Invitrogen, USA) was used for 15 min at room temperature to stain the nuclei.

### 2.5. Digital Analysis of Micrographs

#### 2.5.1. Quantification of Nuclear Deformation

Fluorescence micrographs of the nuclei of cells were analyzed by using the image analysis software ImageJ (NIH)^31^ to determine the “circularity” (Equation 1) of cell nuclei:

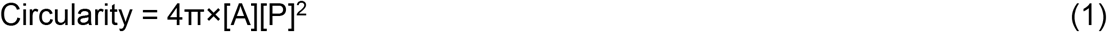

where A is the area and P is the perimeter of the nuclei.

Analyses were performed using 100 cell nuclei per surface. For the auto thresholding of the images, ImageJ was used.^32^

A “Circularity” value of 1 indicates a perfect circle. As the value decreases towards 0, the shape becomes an increasingly elongated polygon.

### 2.6. Quantification of Expression Levels of Proteins Based on the Intensity of ICC Staining

Confocal micrographs of the samples stained with antibodies specific to Lamin A/C and Nesprin-2 were analyzed by using image analysis software Fiji.^33^ Original images were in Red (R), Green (G), Blue (B) format. Image preprocessing was applied to obtain gray scale (8 bit) images. “Lookup table” of Fiji was changed to “HiLo”. Then the contrast was adjusted. Background subtraction was done using “rolling ball algorithm”.^34^ Finally, the intensity of the antibody specific stain was measured and a mean gray value (sum of the gray values of all the pixels in the selection divided by the number of pixels) was obtained for each micrograph.

### 2.7. Statistical analysis

All quantitative data in this study were expressed as mean ± standard deviations with n≥3 unless otherwise stated. Shape analyses were done with 100 cells per surface. Normality test on all collected data was performed by Shapiro-Wilk test. Statistical analysis was performed by one-way or two-way ANOVA (analysis of variance) test followed by Tukey’s test for normally distributed data and Kruskal-Wallis test for non-normally distributed data. Data were presented as symbol plots, where symbols and error bars represent the mean and standard deviation, respectively or presented as box-whisker plots, where boxes represent 25th and 75th percentiles and whiskers represent the values from min to max. The line in the middle of the box is plotted at the median and “+” at the mean. Statistical significance is set at 95% confidence level for all tests (p < 0.05).

## 3. Results and Discussions

Square prism shaped micropillar (8 μm tall, 4×4 μm^2^ area, 4 μm interpillar gap) decorated surfaces (P) were prepared on PDMS using the original wafer as the main template. Unpatterned (smooth) (S) surfaces served as controls. Figure 1 shows a scheme of the workflow in this study. Figure 1A presents SEM micrographs of the micropillar decorated and smooth surfaces and their graphical illustrations (insets). Figure 1B shows the nuclear deformations of three breast cancer cell lines as quantified with the circularity value obtained by Fiji software and representative fluorescence micrographs of their nuclei. Circularity values for the breast cancer cell lines are: 0.77 for benign MCF10A, 0.47 for noninvasive malignant MCF7 and 0.37 for invasive malignant MDAMB231. A lower circularity value implies a high degree of deformation of the nuclei, in other words a soft nucleus. In our previous study, a detailed comparison of these three cell lines were made and it was shown that the nuclei of the more aggressive cancer cells have a higher capability of deformation.^16^ This was also a proof of the ability of metastatic cancer cells migrate through dense ECM by changing the morphology of their nuclei.^35–38^ Figure 1C shows the action mechanisms of the two actin inhibitor drugs Cytochalasin D (CytoD) and Jasplakinolide (Jasp) used in this study. CytodD binds to the positive end of the actin filaments and prevents their polymerization while Jasp binds to negative end and prevents depolymerization which results in aggregation of actin filaments.

**Figure 1:**
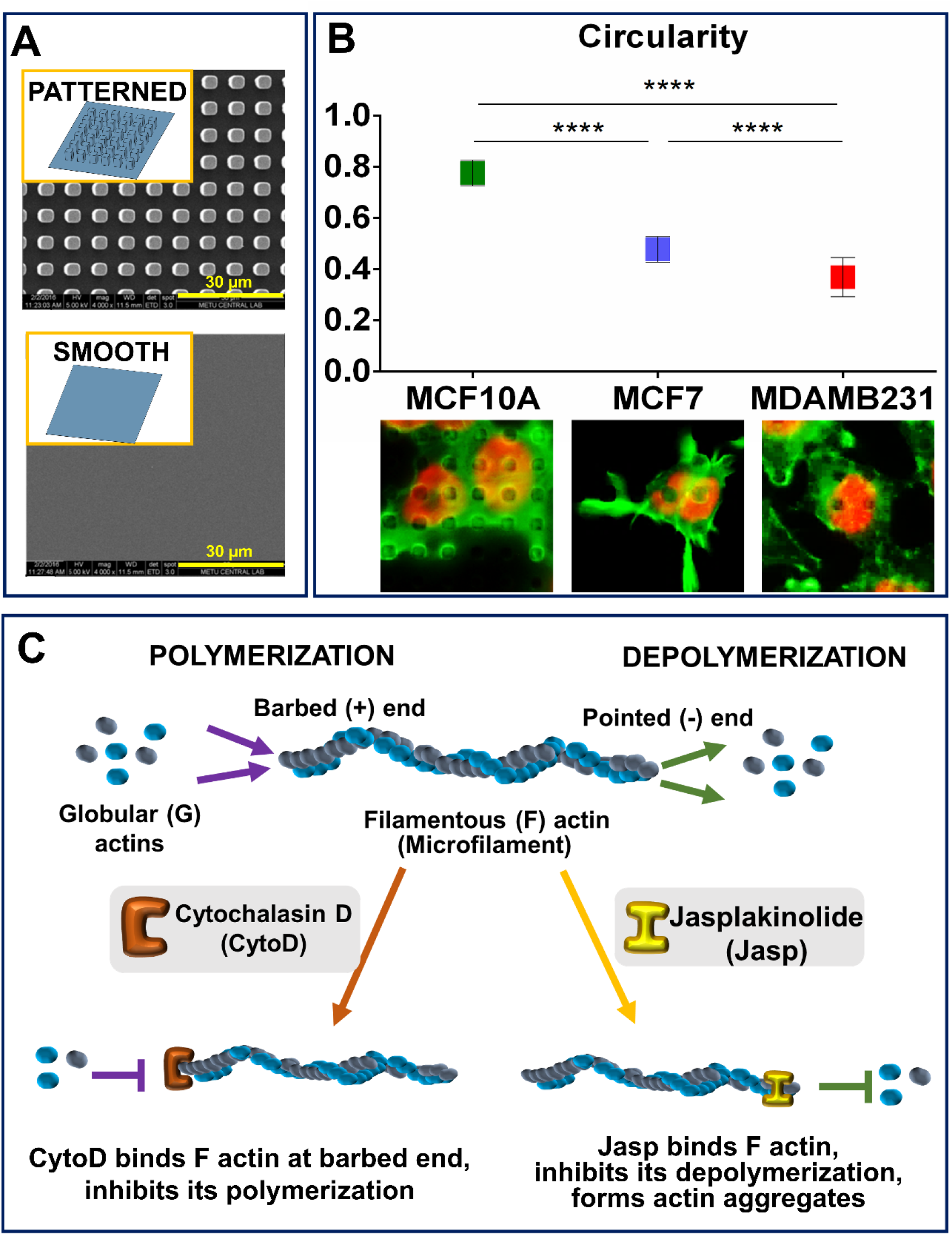
Summary of the nuclear deformability of cells on micropatterned surfaces and the action mechanism of the drugs inhibiting actins. A) SEM of patterned and smooth PMMA surfaces, B) Circularity value of three breast cancer cell lines (The distribution of shape descriptor values was non-normal and statistical analysis was carried out by Kruskal–Wallis non-parametric test, n≥100 cells/surface, *p < 0.005 **p < 0.001, ***p < 0.0005, and ****p < 0.0001. n.s: not significant), C) Acting mechanism of actin inhibitory drugs: CytoD and Jasp.

### 3.1. Drug Testing on Breast Cancer Cells

Drugs inhibiting actin formation were used to study the role of mechanotransduction on the deformability of the cells because it is known that there is a coordination between the mechanosensory elements and force transmission.^39^ External forces applied on a cell are transduced through the actin cytoskeleton to the nucleus and eventually may cause the deformability of the cell and its nucleus.^39^ In the actin inhibiting drug tests, the cells were initially cultured for 24 h in the tissue culture plate, and then treated with the drugs for another 24 h before microscopy. Cytochalasin D used in this study inhibits actin polymerization by binding to the monomers and capping the filaments selectively.^40,41^ Jasplakinolide, on the other hand, stimulates actin polymerization but disrupts F-actin fiber formation by blocking FAK signaling pathway.^42^

Dose-Response curves show that when Cytochalasin D (CytoD) or Jasplakinolide (Jasp) (Supplementary Figure 1) were used at their highest concentration (10 μM and 5 μM, respectively), they decreased metabolic activity of the cells rather than completely stopping proliferation. This decrease was not more than 50% in most of the 3 cell types tested. In the literature, it was also shown that the cytotoxic levels of these drugs were above 100-200 μM against breast cancer cells and the concentrations used in our study were much more below the lethal dose.^42,43^ Thus, the cells treated with the drugs were still showing metabolic activity.

Fluorescence micrographs of the three cancer cells after interaction with the drugs on TCPS are shown in Figure 2. The cells were immunostained with Alexa Fluor 488® labeled Phalloidin (green) for filamentous actin (F-actin) and with DAPI (red) for the nuclei. It is observed that increasing the concentration of drugs led to a progressive decrease in the actin contents as shown by the decrease in the green signal obtained from the actin-specific stain and disorganized and disrupted morphology of the actin cytoskeleton. Similar results were also shown in the literature that increasing CytoD concentration from 10 to 200 μM resulted in progressive shrinking of the cells and reduced F-actin content when metastatic human cancer cell lines were used.^42^

The highest concentrations of the both drugs (10 μM for CytoD and 5 μM for Jasp) showed a distinct effect on the actins of the cells (Figure 2), so these two highest drug doses were selected to use in the nuclear deformation analysis.

**Figure 2:**
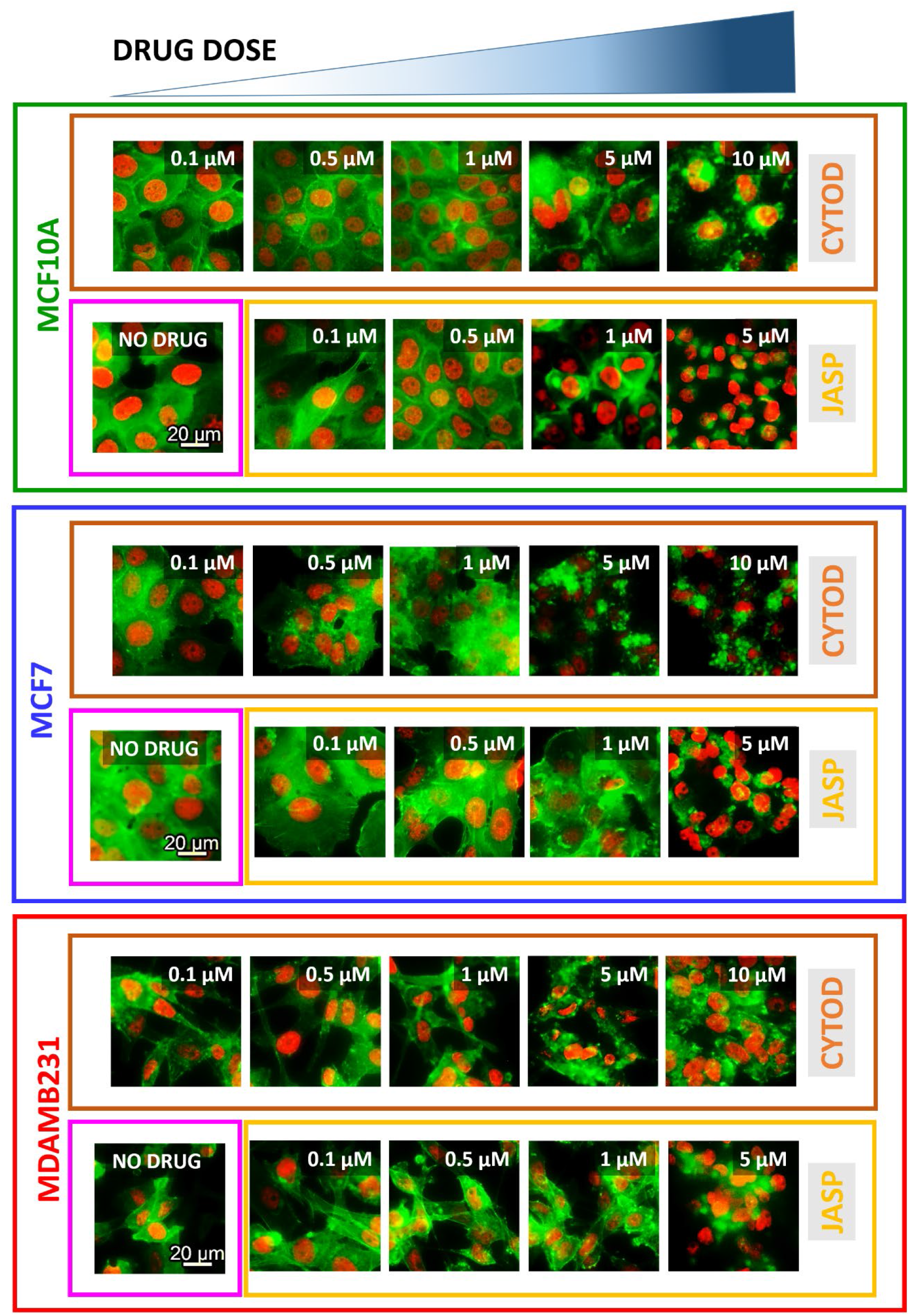
Fluorescence micrographs on the effect of concentration of the drugs (Cytochalasin D (CytoD) and Jasplakinolide (Jasp)) on actin cytoskeleton formation (Green: Alexa Fluor 488 Phalloidin) of the three breast cancer cells on TCPS. (Nucleus: Red (DAPI), Scale bar: 20 μm).

Fluorescence micrographs of the nuclei of the cells on micropatterned PMMA surfaces are presented in Figure 3. Upper two row micrographs show cells after 24 and 48 h drug-free incubation. The lower rows show the cells after drug treatment (drug free 24 h incubation followed by 24 h with the drug). After 24 h without the drugs the nuclei of benign cells (MCF10A) were not deformed whereas nuclei of malignant cells MCF7 and MDAMB231 were highly deformed. After 48 h of drug-free incubation, nuclei of all the cell types, regardless of being benign or malignant, were deformed. Thus, the malignant cell types were deformed earlier than the benign ones on micropatterned surfaces.

**Figure 3:**
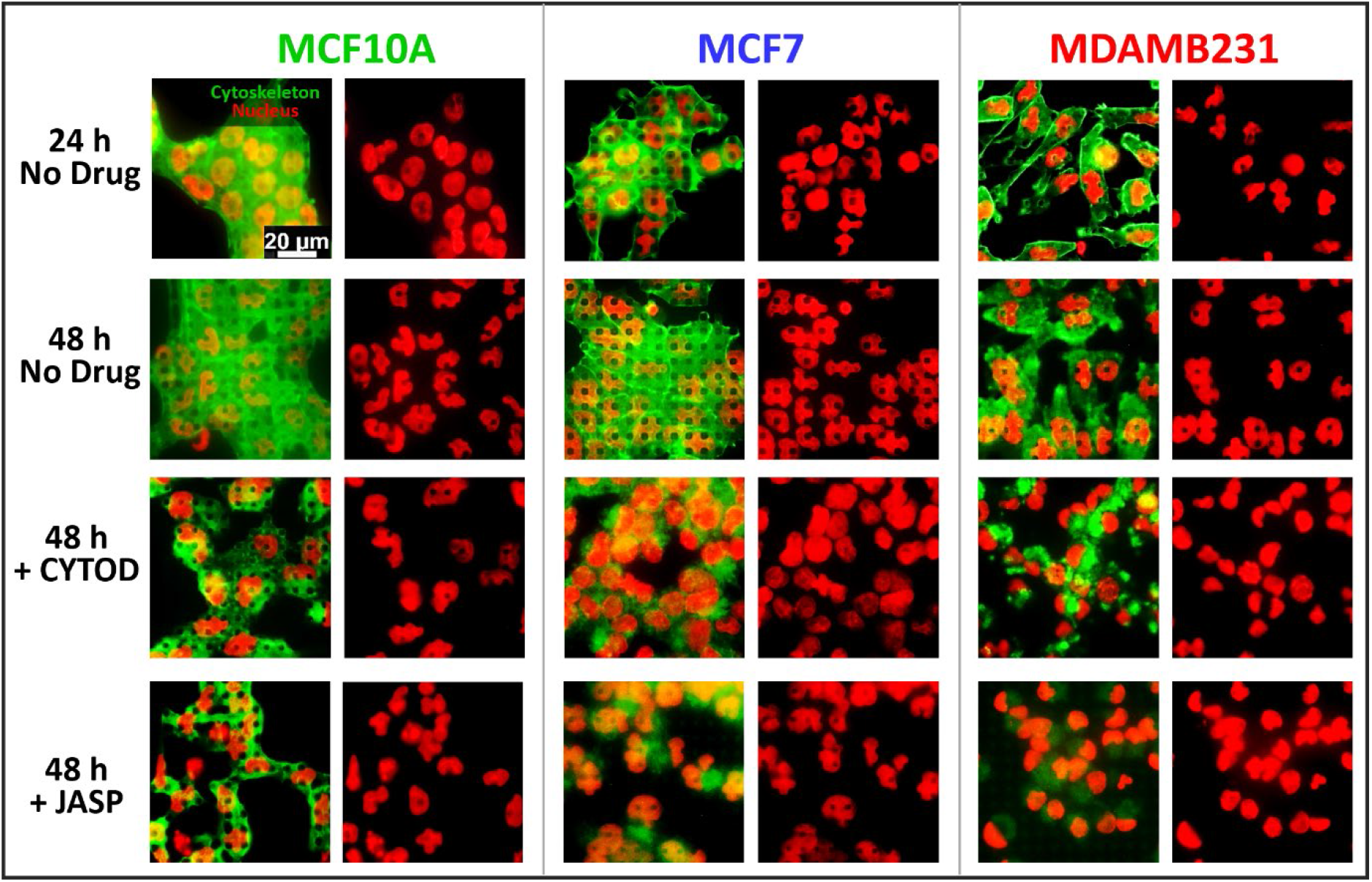
Fluorescence micrographs show the effect of drugs (CytoD, 10 μM and Jasp, 5 μM) on actin cytoskeleton by the decrease in the green signal of actin specific stain (Alexa Fluor 488 Phalloidin) and nuclear deformation (DAPI, Red) of cells on micropatterned PMMA surfaces after 24 h and 48 h without drug (upper two rows) and 48 h after drug treatment (bottom two rows) (Scale bar: 20 μm).

However, after 24 h incubation with the CytoD, the nuclei of benign cells were less deformed (MCF10A) while malign cells were not deformed at all (MCF7 and MDAMB231). With Jasp, on the other hand, the nuclear deformation of the cells was decreased MCF10A and MCF7 but this decrease was significant only for MDAMB231. Thus, CytoD was more effective than Jasp in decreasing the deformability of the cells. It is also observed in the micrographs that the drugs disrupted the actin filaments of the cells and simultaneously the nuclear deformations were decreased or lost. This shows that nuclear deformability is directly related to the organization of actins. Studies showed that loss or disruption of actins caused loss of actin-myosin contractility and undeformed, force-free state of cells.^42,44^ It was also stated in a review that the actomyosin network is responsible for contractility and force generation in the cell and coordination between the mechanosensors (integrins) and response elements (actin cytoskeleton) guides force transduction and the deformability of the cell.^39^ This supports the observation of loss of deformability after disruption of actin filaments in our study.

### 3.2. Quantification of Nuclear Deformation of Cells

Extent of deformation of the cell nuclei was quantified by digital analysis of the fluorescence micrographs using Fiji software. As a measure of deformation, certain dimensional properties of the cells were employed. Circularity is one such parameter and is a measure of how close a cell nucleus is to a perfect circle. A circularity of 1 indicates a perfect circle whereas a zero is a substantially elongated shape. The equation defining the circularity was presented in Section 2.5.1.

Circularity values of the nuclei of the three cells on micropatterned PMMA surfaces before and after the treatment with CytoD and Jasp are presented in Figure 4.

**Figure 4:**
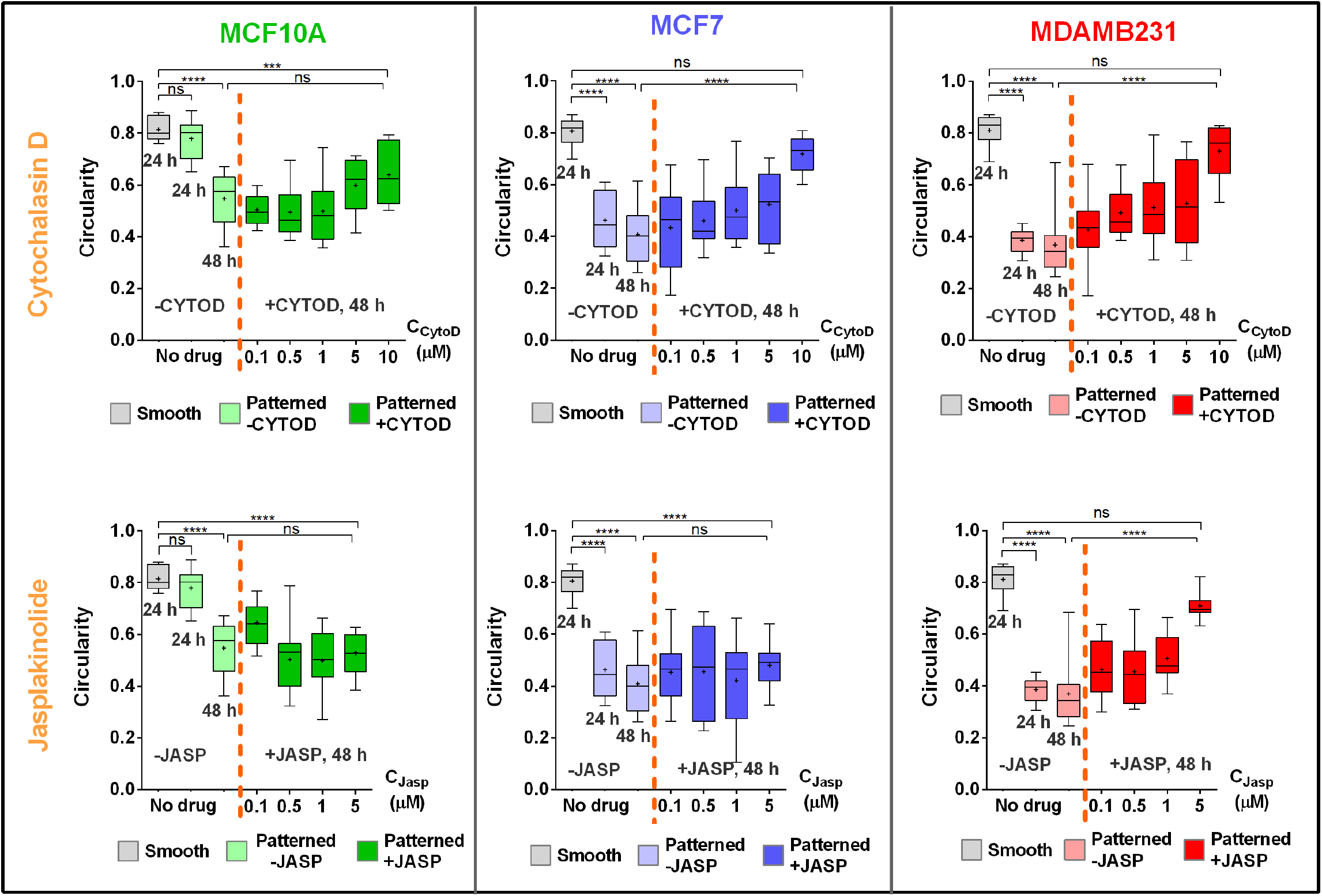
Quantification of nuclear deformation by the circularity value calculated from fluorescence micrographs of the three breast cancer cells before and after CytoD (10 μM) and Jasp (5 μM) treatment (Statistical analysis was carried out by using one-way ANOVA, n≥100 cells/surface, *p < 0.05, **p < 0.01, ***p < 0.001, ****p < 0.0001. n.s: not significant).

On smooth (unpatterned) surfaces, circularity values for all cells were around 0.8 which is almost a perfect circle value indicating that there is no nuclear deformation.

On micropatterned surfaces, after 24 h, circularity values were 0.78, 0.46 and 0.39 for MCF10A, MCF7 and MDAMB231 cells, respectively. This means that the nuclei of benign MCF10A cells had almost no conformational change whereas nuclei of the cancer cells were severely deformed. Increase in the nuclear deformation by the increase in the malignancy of cells also correlates well with the overall stiffness of the cells and their nuclei obtained in a study.^45^ Young’s Modulus values were found to be 0.7, 0.5 and 0.3 kPa for MCF10A, MCF7 and MDAMB231 cells, respectively, supporting the studies reporting that the softest cancer cells were also most deformed on the micropatterned surfaces.^10,13,38^ After 48 h, nuclei of all cells were deformed somewhat more; circularity values were now 0.55, 0.41 and 0.37 for MCF10A, MCF7and MDAMB231, respectively. Results show that eventually, all the cells are deformed but the deformation was slower for benign (MCF10A) cells whereas nuclei of invasive malignant (MDAMB231) cell deformed at a much higher rate. These results were very similar to the results reported earlier on the discrimination of breast cancer cells by their malignancies on micropatterned surfaces.^16^

In the tests for drugs acting on actin polymerization and organization, the cells were treated with CytoD and Jasp and then nuclear deformations of the cells were quantified as above:

After the treatment of cells with CytoD concentrations in the range 0.1-10 μM, circularity values of all cells increased with an increase in the drug concentration and approached 1.0 even though they were on micropatterned surfaces. The circularity values after the treatment with the highest concentration (10 μM) of CytoD were 0.64, 0.72 and 0.73 for MCF10A, MCF7 and MDAMB231, respectively. Benign (MCF10A) cells were not affected by the drug treatment as much as malignant cells (MCF7 and MDAMB231). Malignant cells lost deformability completely upon treatment with the high dose of drug.

When the same test was performed with Jasp (0.1-5 μM), only the nuclei of the most malignant cell line, MDAMB231, were affected with a significant increase in the circularity (decrease in the deformation). The circularity values of the three cells after the highest concentration of Jasp (5 μM) were 0.53, 0.48 and 0.71 for MCF10A, MCF7 and MDAMB231, respectively.

It is known that actin cytoskeleton of cells not only provides structural support but also defines the mechanical properties of the cells. Actin network is involved in the maintenance of nuclear stiffness and actin filaments are needed for the cell relaxation.^29,30^ Thus, in our study, after the disruption of the actin cytoskeleton by the drugs, cells lost their actomyosin contractility and this resulted in the changes observed in their deformability. It can be concluded that there is a direct connection between nuclear deformation and actin filaments of the cells.

### 3.3. Expression of Lamin A/C and Nesprin-2 Before and After Drug Treatment

Lamin A/C (nuclear lamina component) and Nesprin-2 (one of the components of the LINC complex) are known to be downregulated in human breast cancer cells.^46^ They play crucial roles in nuclear membrane organization and mechanical stiffness and their downregulation leads to decrease of nuclear and cellular rigidity and increase of tissue fluidity which is a crucial property of malignant cancer cells.^46^ Expression levels of these proteins decrease substantially in breast cancer cells and this results in an increase in the nuclear deformability of these malignant cells. We chose malignant MCF7 as a model for immunocytochemistry (ICCI staining and expression of Lamin A/C and Nesprin-2 were studied with this approach. The cells were seeded on micropatterned (P) and smooth (S) PMMA films and tissue culture plates (TCPS). Their cytoskeletal and nuclear morphologies were recorded with Confocal Laser Scanning Microscopy (CLSM) after 24 h incubation in culture medium followed by another 24 h with the drug CytoD and the micrographs on micropatterned surfaces are presented in Figures 5A and 5B whereas those on smooth and TCPS surfaces are presented in Supplementary Figures S2 and S3. Protein expression levels were calculated from the intensity of the stains in the confocal micrographs (Figures 5C and D).

**Figure 5:**
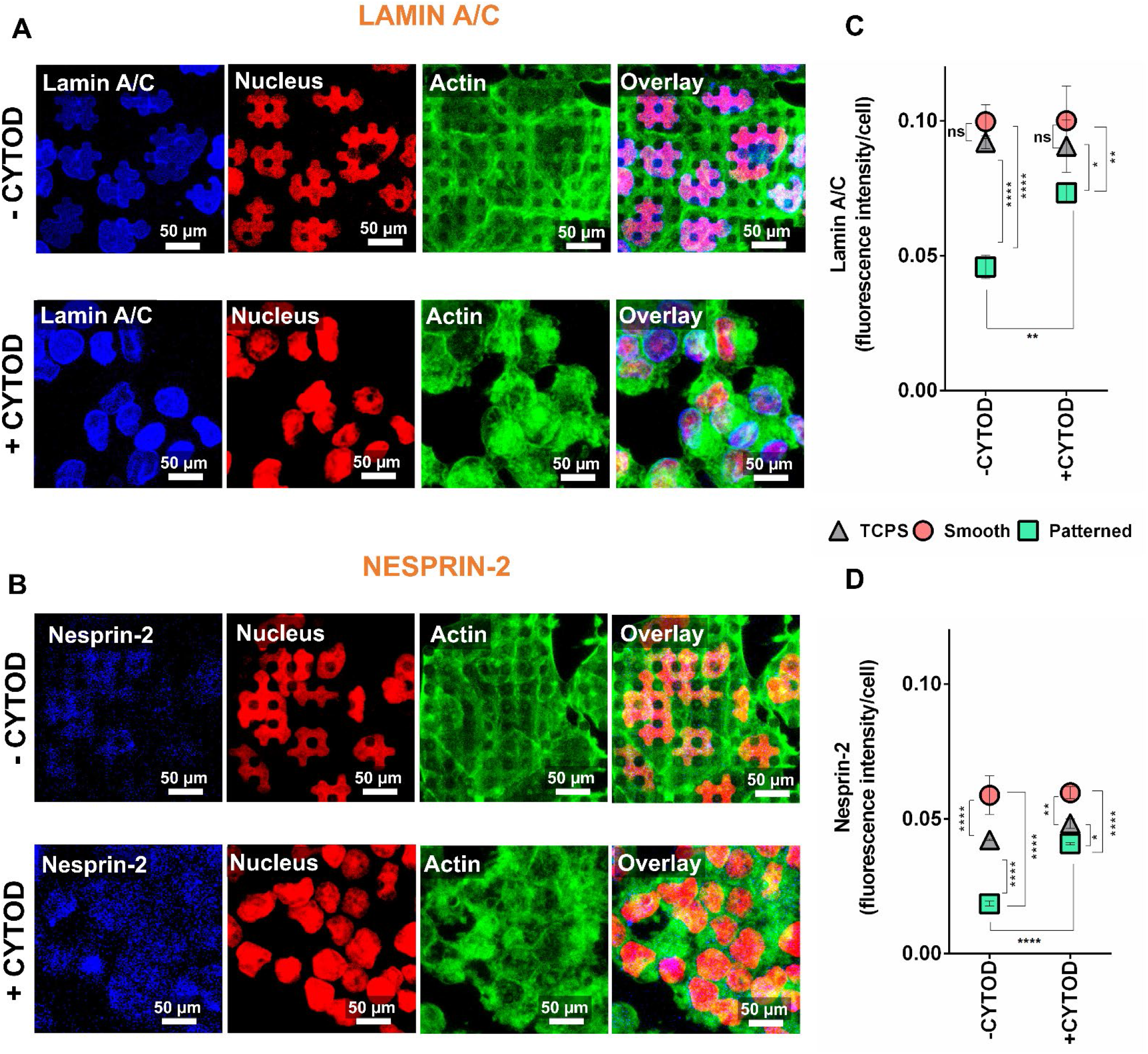
ICC staining of MCF7 malignant breast cancer cells with Lamin A/C and Nesprin-2 specific antibodies. CLSM micrographs of cells on micropatterned surfaces stained for **A)** Lamin A/C and **B)** Nesprin-2 (blue: Anti Lamin A or Anti-Nesprin-2), nucleus (red: TOPRO-3) and actin filaments (green: Alexa Fluor 532 Phalloidin) (X40 objective, Scale bars: 50 μm). Quantification of expression levels of **C)** Lamin A/C and **D)** Nesprin-2 before and after drug treatment as determined from their fluorescence intensities on micropatterned, smooth and TCPS surfaces. Fluorescence intensities were calculated using ImageJ (NIH) software. Results were given as dividing signal intensities by the number of cells counted from the same surface (Two-way ANOVA followed by Sidak’s multiple comparison test, * p<0.05, ** p<0.01, *** p<0.001, p<0.0001****, n=3).

Micrographs before drug addition show that nuclei of MCF7 cells were deformed on P4G4 micropatterned surfaces (Figures 5A and 5B, first rows).

After the addition of drug CytoD, the actin cytoskeletons of the cells were disrupted as shown by the actin specific stain (Green: Alexa Fluor 532 Phalloidin) and the nuclear deformability of the cells were lost on micropatterned surfaces (Figures 5A and 5B, second rows). However, signal intensities of both Lamin A/C and Nesprin-2 were increased (Blue: Anti Lamin A or Anti-Nesprin-2) (Figures 5C and 5D) as was measured and quantified with ImageJ (NIH) software. Results are optimized by the cell number; signal intensities were divided by the number of cells on the same surface. The results show that after drug treatment and simultaneously with loss of nuclear deformability signal intensities of Lamin A/C and Nesprin-2 were increased on patterned surfaces (Figures 5C and 5D). This indicates that the expression levels of these two proteins are increased. Lamin A/C and Nesprin-2 are in a direct contact with actin filaments of the cell. CytoD caused the disruption of actin filament formation by inhibiting their polymerization, and therefore, cells lost actomyosin contractility, and therefore, deformability. In a study, when dendritic cells were forced to pass through micrometric constrictions, it was observed that actin polymerization around the nucleus was required for the cell to pass through small constrictions since they these fibers are involved in actomyosin contractility and force transduction in the cell which start from the cell membrane and end up with the nucleus.^47^ Moreover, it was shown that intermediate lamin A/C and B proteins form a rigid structure at the inner nuclear membrane and cells need to express lamin gene product less to be able to migrate.^47^ This is why deformable and softer cancer cells show lower Lamin A/C protein expressions.

In summary, our results show that after the disruption of actin filaments, the breast cancer cells lost their capability to deform and expression of Lamin A/C and Nesprin-2 proteins increased. Based on these results and the literature data, we suggest that the migrating and metastatic cells need a low level of these two proteins to become more flexible and deformable. Moreover, since the actin disruption cause the decrease in nuclear deformation, cells do not require low expression levels of the two proteins (which were considered as barriers limiting the nuclear deformability). Thus, their expression increased after the loss of nuclear deformation.

On TCPS and smooth surfaces, on the other hand, intensities of the proteins did not change significantly, and nuclear deformation was not observed.

## 4. Conclusion

Cancerous and healthy cells placed on micropatterned surfaces show differences in various morphological and biomechanical properties. Deformability is a property specific for cancerous or diseased cells and it is important to study the underlying mechanism. Besides, deformability carries hints about the metastatic capability and the malignancy of the cells. In this study, three breast cancer cell lines with different malignancy levels were cultured on micropatterned surfaces. Nuclear deformation levels, expression levels of the mechanotransduction proteins and F-actin contents of these cells were measured before and after the use of actin inhibiting drugs. As a result, actin inhibiting drugs revealed that actin filaments, and Lamin A/C and Nesprin-2 proteins have distinct role in the force transmission and nuclear deformation. It is the first time in the literature that we show the nuclear deformability of the malignant breast cancer cells can be prevented by the disruption of these proteins and this can be considered as a crucial data in the recovery of metastatic behavior of cancer cells.

## Acknowledgements

EA and VH acknowledge the Ministry of Development of Turkey, METU BAP-01-08-2013-003 and BAP-08-11-DPT2011K120350, METU BIOMATEN and TUBITAK 2211-C scholarship. UD would like to acknowledge NIH R15HL115556. This material is based in part upon work supported by the National Science Foundation under NSF CAREER Award Number 1461602 and NSF EAGER 1547791. UD would also like to acknowledge Canary Foundation seed grant, NCI Center for Cancer Nanotechnology Excellence for Translational Diagnostics (NIH NCI U54CA199075).

## Supplementary Figures

**Supplementary Figure S1:**
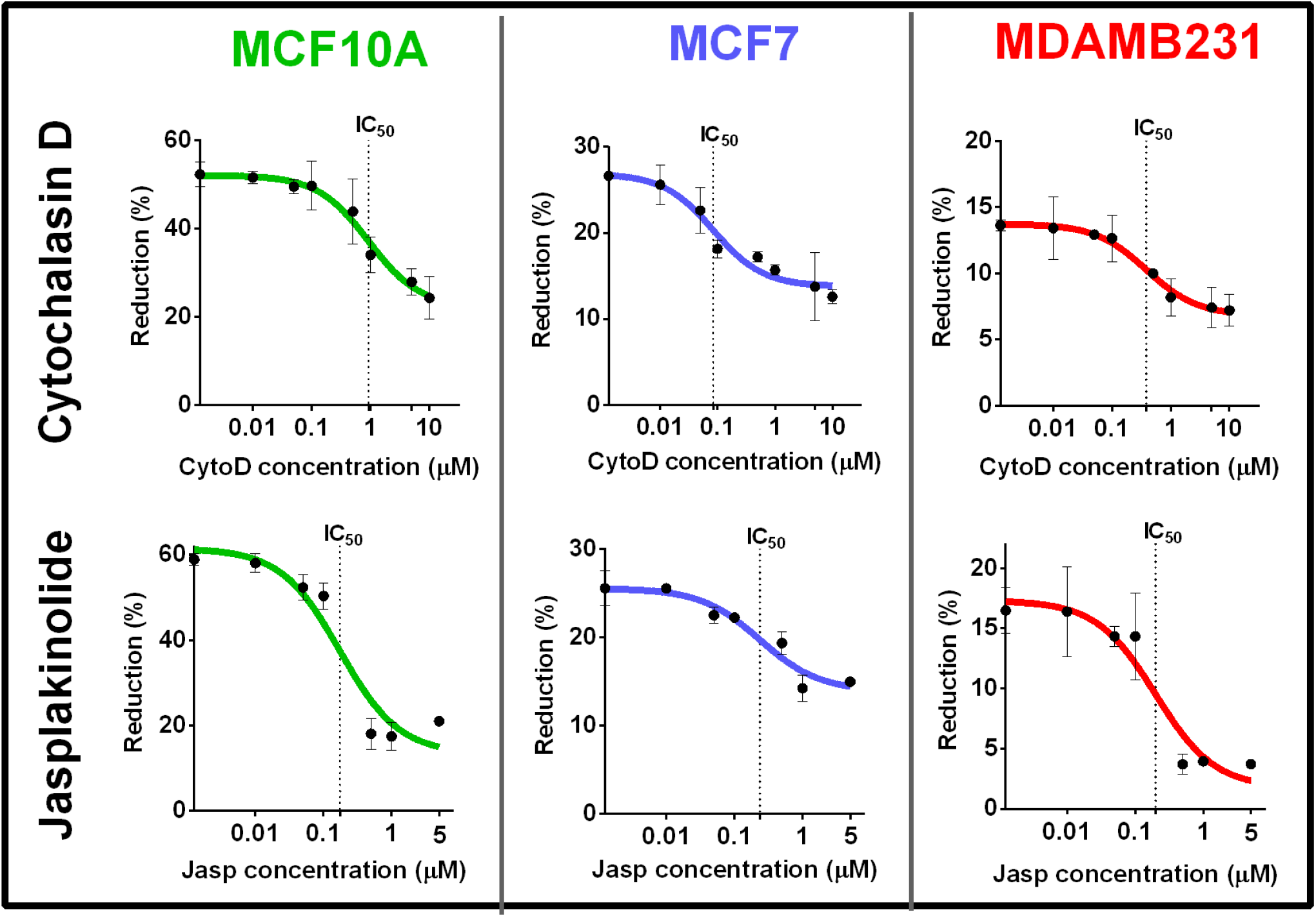
Drug dose-response curves of Cytochalasin D (CytoD) and Jasplakinolide (Jasp) on 3 cell lines. Serial dilution of drug concentrations on TCPS are shown by using reduction (%) calculated from Alamar Blue Assay. Dose axis is logarithmic. IC_50_ indicated the drug dose needed to decrease the drug activity to its 50%.

**Supplementary Figure S2:**
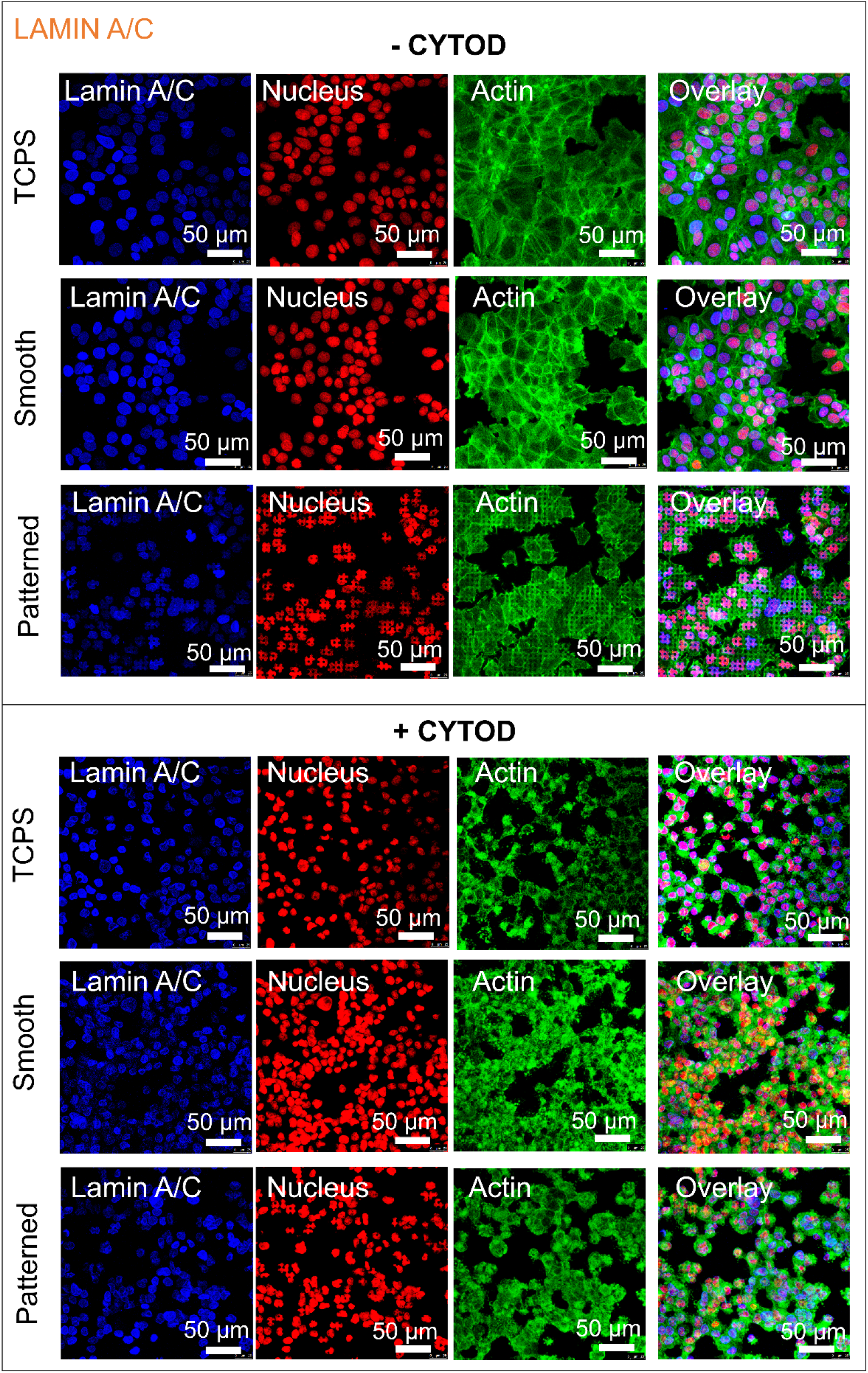
ICC staining of MCF7 malignant breast cancer cells with Lamin A/C specific antibody before and after CytoD treatment on micropatterned, smooth and TCPS surfaces. Blue: Anti Lamin A/C, Red: TOPRO-3 (Nucleus), and Green: Alexa Fluor 532 Phalloidin (Actin filaments). (X40 objective, Scale bars: 50 μm).

**Supplementary Figure S3:**
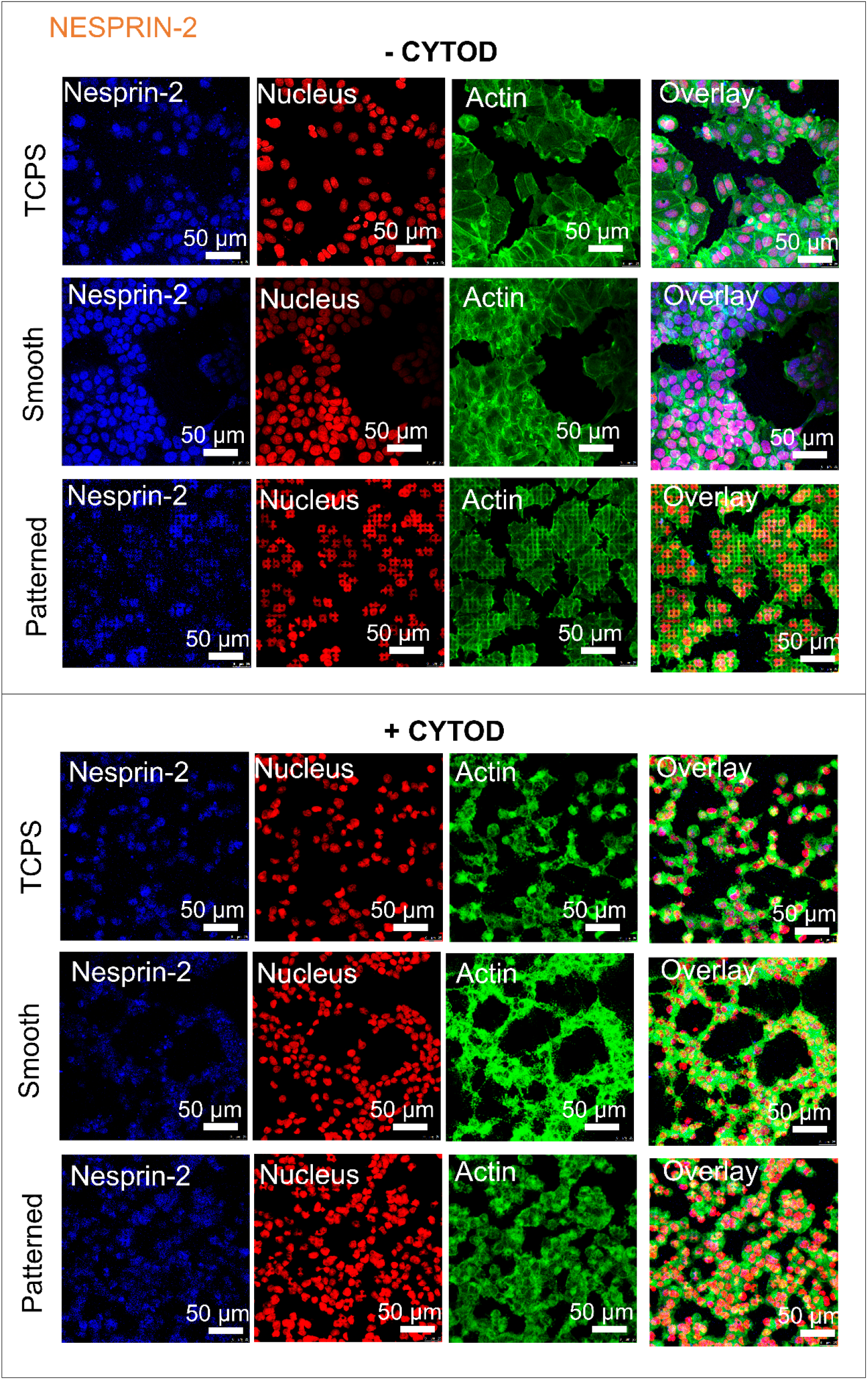
ICC staining of MCF7 malignant breast cancer cells with Nesprin-2 specific antibody before and after CytoD treatment on micropatterned, smooth and TCPS surfaces. Blue: Anti-Nesprin-2, Red: TOPRO-3 (Nucleus), and Green: Alexa Fluor 532 Phalloidin (Actin filaments). (X40 objective, Scale bars: 50 μm).

